# Multi-objective prioritisation of candidate epitopes for diagnostic test development

**DOI:** 10.1101/2021.09.17.460787

**Authors:** Roman Cerny, Jodie Ashford, João Reis-Cunha, Felipe Campelo

## Abstract

**Background:** The development of peptide-based diagnostic tests requires the identification of epitopes that are at the same time highly immunogenic and, ideally, unique to the pathogen of interest, to minimise the chances of cross-reactivity. Existing computational pipelines for the prediction of linear B-cell epitopes tend to focus exclusively on the first objective, leaving considerations of cross-reactivity to later stages of test development.

**Results:** We present a multi-objective approach to the prioritisation of candidate epitopes for experimental validation, in the context of diagnostic test development. The dual objectives of uniqueness (measured as dissimilarity from known epitope sequences from other pathogens) and predicted immunogenicity (measured as the probability score returned by the prediction model) are considered simultaneously. Validation was performed using data from three distinct pathogens (namely the nematode *Onchocerca volvulus*, the Epstein-Barr Virus and the Hepatitis C Virus), with predictions derived using an organism-specific prediction approach. The multi-objective rankings returned sets of non-dominated solutions as potential targets for the development of diagnostic tests with lower probability of false positives due to cross-reactivity.

**Conclusions:** The application of the proposed approach to three test pathogens led to the identification of 20 new potential epitopes, with both high probability and a high degree of exclusivity to the target organisms. The results indicate the potential of the proposed approach to provide enhanced filtering and ranking of potential candidates, highlighting potential cross-reactivities and including this information into the test development process right from the target identification and prioritisation step.

## Background

Computational prioritisation of peptide targets to be used in antibody-derived serologic diagnostic tests is commonly based on three main determinants: Predicted antigenicity, sequence conservation among isolates of the pathogen of interest, and sequence dissimilarity to known epitopes from other pathogens (Can et al., 2020; Reis-Cunha et al., 2014; Gourlay et al., 2017). The combination of these criteria results in a multi-criterion decision-making problem, in which the trade-offs between the expected immunogenicity of a given peptide and its uniqueness to the pathogen of interest must be simultaneously considered.

Computational methods, including machine learning (ML) algorithms, are now commonly used strategies for B-cell epitope prediction. These methods help to overcome some of the limitations imposed by traditional experimental epitope identification techniques (Yang and Yu, 2009; Haste Andersen et al., 2006; Yasser and Honavar, 2010). Several ML algorithms and pipelines have been used for linear B-cell epitope prediction with varying success, including Random Forest classifiers (Jespersen et al., 2017; Saravanan and Gautham, 2015; Yasser and Honavar, 2014), Gradient Boosting (Manavalan et al., 2018), Support Vector Machines (Chen et al., 2007; EL-Manzalawy et al., 2008; Sweredoski and Baldi, 2009; Wee et al., 2010; Wang et al., 2011; Gao et al., 2012; Lin et al., 2013; Shen et al., 2015; Yao et al., 2012; Singh et al., 2013) and Artificial Neural Networks (Saha and Raghava, 2006; Sher et al., 2017; Lian et al., 2015; Collatz et al., 2021). Though, generally, these ML methods outperform traditional experimental and older computational methods, like amino acid propensity scales (Blythe and Flower, 2005; Giacò et al., 2012; Sanchez-Trincado et al., 2017), these current methods still exhibit relatively low prediction performance (Yang and Yu, 2009; Ashford et al., 2021).

While these performance issues may be partially explained by the intrinsic hardness of the epitope prediction problem and the low information content of commonly used predictive features, a significant contributor may be the selection of the training data itself. Currently, most sequence-based epitope predictors are trained as generalist models, based on large, heterogeneous training sets containing labelled sequences from a wide variety of organisms (Ashford et al., 2021). However, recently, epitope prediction works have emerged that use taxon- or organism-specific data sets for training (Ong et al., 2021; Ashford et al., 2021), with substantially improved prediction performance when compared to generalist training.

Antigenicity estimation by B-cell epitope predictors allows the identification of protein regions with a higher chance to be recognized by antibodies. This is a fundamental step in the development of vaccines, serologic diagnostic assays and other immunodiagnostic tools (Shirai et al., 2014; Soria-Guerra et al., 2015). Selected proteins with a high density of B-cell epitopes can be recombinantly expressed in bacteria or yeast (Nascimento and Leite, 2012), while short stretches of highly antigenic amino acids can be synthesised as peptides, to be used in diagnostic tests (Kamath et al., 2020; Suárez-Fariñas et al., 2021).

Besides antigenicity, sequence conservation and dissimilarity are also crucial for the development of accurate serologic diagnostic tests. The potential sensitivity of the target can be improved by selecting antigenic regions that are conserved in isolates of the same pathogen, obtained from different geographic regions and/or in different time spans (Luo et al., 2015; Mosa, 2020). Similarly, the potential positive predictive value of a diagnostic target can be improved by removing sequences that are conserved in other pathogens that could result in cross-reactivity, as well as sequences that are present in the host (de Oliveira Mendes et al., 2013; Reis-Cunha et al., 2014). These considerations are not incorporated in current epitope predictor tools, where researchers usually run both steps sequentially and independently. Hence, a multi-objective framework combining epitope prediction with sequence conservation/uniqueness could potentially facilitate diagnostic target selection, as well as provide some insight into which pathogens could result in potential cross-reactivity for serologic tests at an early stage of development.

This work presents a prioritisation approach for antibody-based diagnostic test development, based on the multi-objective ranking of targets generated by any epitope prediction tool that outputs probabilities. Two objectives are considered at this stage: the predicted probability of a given peptide representing a linear epitope, which should be maximised; and the similarity between the candidate peptide and known epitopes from other pathogens, which should be minimised in the case of diagnostic test development. To generate priority rankings that consider both criteria simultaneously, non-dominated sorting (Deb et al., 2002; Goulart and Campelo, 2016) is used as the prioritisation approach. Although this work demonstrates this concept using only the two objectives outlined above, it can be easily expanded to include, e.g., conservation among isolates of the pathogen of interest or any other aspect that the user may want to optimise when prioritising targets for experimental investigation.

To illustrate the proposed approach, predictions generated for three distinct organisms were used. These pathogens include the nematode *Onchocerca volvulus* (taxonomy ID: 6282) and two viruses (Epstein-Barr Virus - taxonomy ID: 10376; Hepatitis C Virus - taxonomy ID: 11102). The predicted probabilities used (objective 1, to be maximised) were those output by the organism-specific models developed in an earlier work (Ashford et al., 2021), for a hold-out subset of proteins from each organism. The similarity score of a given target (objective 2, to be minimised) was calculated as the largest alignment score value between the predicted sequence and and all known epitopes documented in the *Immune Epitope Data Base* (IEDB) (Vita et al., 2019).

## Results

The organism-specific models described by Ashford *et al*. (Ashford et al., 2021) were employed to generate predictions for the proteins that were allocated to the hold-out set of each pathogen. Continuous peptides formed of at least 8 sequential positions with a predicted probability greater than 0.5 were marked as positive predictions. Out of those, the ones having strictly more than 50% of their positions not having a known class (based on the available IEDB data) were selected as potential *new* epitopes. The *Probability* values for these peptides was set as the mean of their position-wise probability scores, and their *Uniqueness* scores were calculated based on the methodology outlined in the “Methods” section, using gap opening and extension penalties of 5 and 2, respectively.

For the prioritisation process, constraints were imposed both on minimal probability and minimal uniqueness. These values were set as *p*_*min*_ = 0.75 and *u*_*min*_. Figure 1 illustrates the distribution of predicted peptides in the decision space, highlighting the three first non-dominated fronts and the feasible region.

**Table 1:**
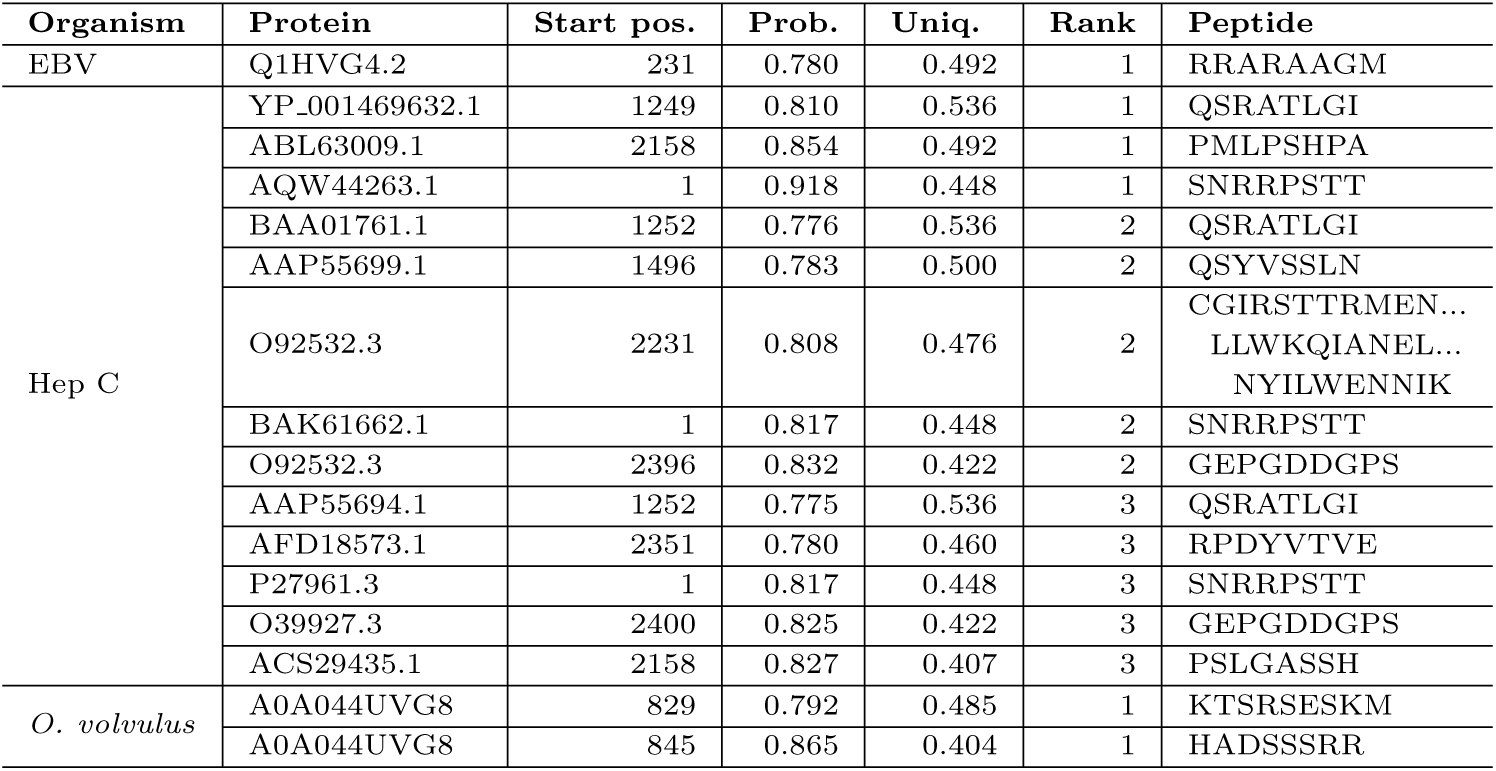
Predicted feasible (probability greater than 0.75, uniqueness greater than 0.3) and new (fewer than 50% of positions having a known label on IEDB) linear B-cell epitopes for the test pathogens, up to Rank 3.

**Figure 1:**
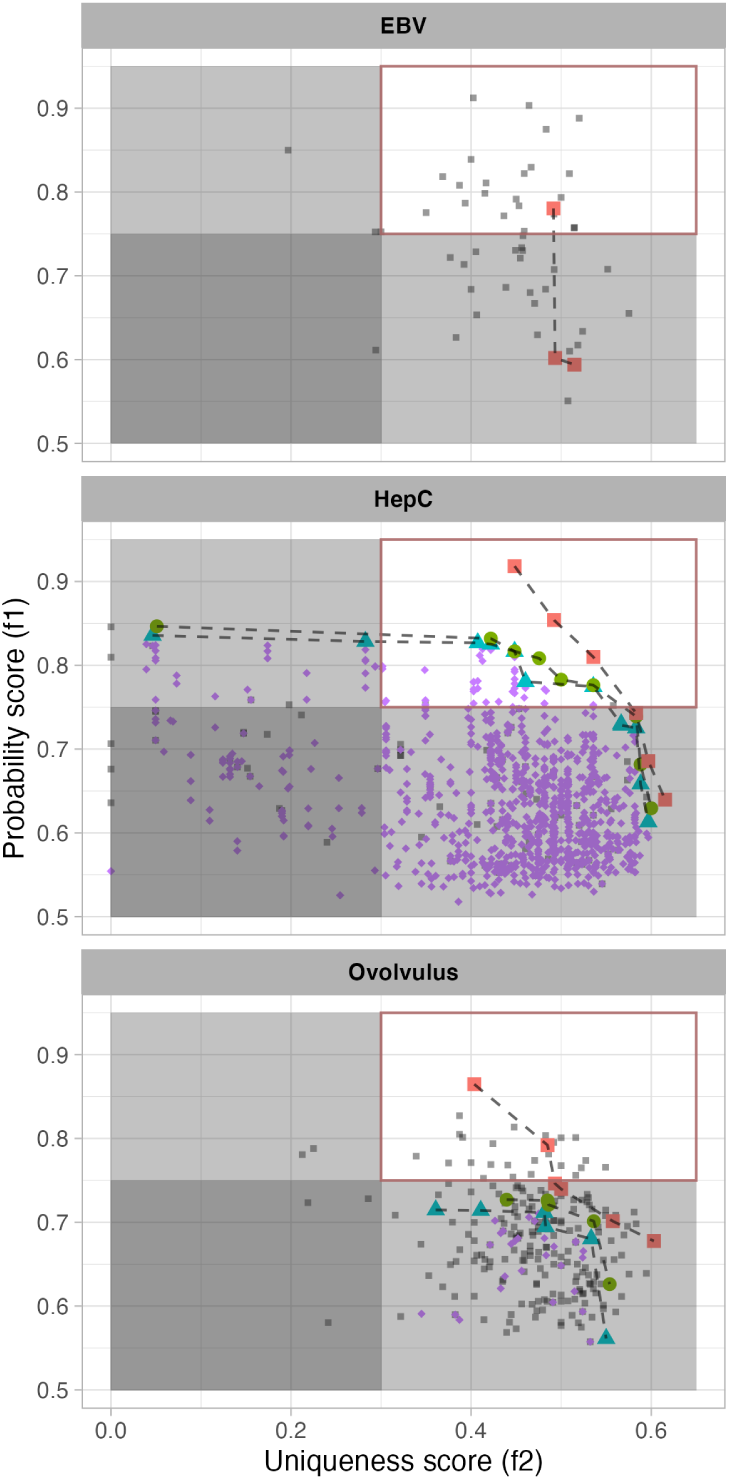
Multi-objective prioritisation of predicted epitopes, for the Epstein-Barr Virus, Hepatitis C and *Onchocerca volvulus*. The first three fronts extracted using non-dominated sorting on the peptides that would represent **new** epitopes (i.e., those for which no labels were available for more than 50% of their length) are highlighted by connected points (Rank 1: red squares; Rank 2: green circles; Rank 3: Blue triangles). Potential **new** epitopes with Rank 4 and beyond are shown as purple diamonds, and all other predicted peptides (i.e., the ones that correspond to known epitopes from the target pathogen) are shown as light gray, smaller squares. The feasible region of the decision space (assuming a problem definition with *p*_*min*_ = 0.75 and *u*_*min*_ = 0.3 in problem formulation) is highlighted at the top-right of each panel. Unfeasible areas are shaded in gray.

The potential advantages of performing candidate prioritisation using the proposed multi-objective approach are particularly clear in the case of the Hepatitis C predictions. For this pathogen, we can observe a large number of high-probability (above 0.8) candidates with uniqueness scores approaching zero, indicating peptides with almost-perfect local alignment to known epitopes from other pathogens. These candidates represent peptides that, although probably correctly identified as epitopes, would represent potentially poor candidates for diagnostic test development, as the likelihood of cross-reactivity with other pathogens is high.

Another interesting aspect that can be observed in Figure 1 is the fact that very few potential new epitopes were predicted for the Epstein-Barr virus (only three under the criteria outlined above, out of 53 predicted peptides). This may indicate that most of the linear B-cell epitopes from the proteins that have already been experimentally investigated for this particular pathogen may have already been identified. Investigation of other proteins from this pathogen (including the unlabelled portions of the training set used for training the organism-specific model, as well as proteins that are not referred by any entry in the IEDB) would be informative, but fall outside the scope of the present work.

## Discussion

In the present work, we have used a multi-objective approach to prioritize epitope candidates for the development of serologic diagnostic tests. The need for multi-objective optimisation concepts emerge whenever there is need to simultaneously consider multiple, potentially conflicting objectives, which is a common occurrence in several branches of science and engineering (Goulart and Campelo, 2016) including bioinformatics (Handl et al., 2007). With the exception of very particular cases, the objectives being optimised exhibit conflicting characteristics, such that it is impossible to simultaneously optimise both - leading to the existence of *trade-offs* that need to be considered when deciding which solutions should be selected. This is the case of antigenicity prediction and sequence uniqueness.

By combining organism-specific epitope predictions and antigen uniqueness, we were able to identify 13, 2 and 1 highly promising epitopes for the diagnostics of respectively Hepatitis C Virus, *Onchocerca volvulus* and Epstein-Barr Virus. The uniqueness of each of these candidates was evaluated by comparison with all other out-of-organism epitopes in IEDB (Vita et al., 2019). Hence, there is a higher chance that they are exclusive to their respective pathogens. This contrasts with approaches that compare potential targets only with a restrict data set of potentially known cross-reacting pathogens in downstream analysis (de Oliveira Mendes et al., 2013; Reis-Cunha et al., 2014). As infections with different pathogens is common in humans, and immunological memory can be long-lasting for B-cells (Cancro and Tomayko, 2021), the proposed approach can mitigate the chance of false positives, even with pathogens where a cross-reaction with the species of interest was previously not known. This approach, however, may be overly conservative, as it considers known epitopes from pathogens with little to no geographical overlap with the target organism, or which cause clinically distinct manifestations, as contributing equally to the uniqueness score. Future versions will refine the method by allowing users to define different weights for existing epitopes based, e.g., on geographic distribution or clinical aspects of disease, providing more flexibility to the prioritisation routines.

Even though the current project was focused on diagnostic target selection, the framework can be easily adapted to other objectives, such as the development of pan-vaccines (Darricarrère et al., 2018; Swenson et al., 2005). This can be achieved by using sequence conservation among different pathogens instead of uniqueness as a second objective, prioritizing epitopes that are identical or highly similar in all the species of interest.

## Conclusions

To improve *in silico* diagnostic target selection, we have developed an multi-objective approach based on predicted antigenicity and uniqueness, leading to the identification of 20 new potential epitopes with high probability and a high degree of exclusivity to the organisms, for important human pathogens. This approach could be easily adapted to address different but relevant challenges, such as pan-vaccine development or any other task where multiple conflicting objectives need to be simultaneously considered early in the development cycle.

Results obtained for three test pathogens indicate the potential of the proposed approach to provide enhanced filtering and ranking of potential candidates, by highlighting potential cross-reactivities (using sequence uniqueness as a proxy) and including this into the test development process right from the target identification and prioritisation step.

## Methods

### Multi-objective ranking

In this work, we are interested in simultaneously maximising two objectives: the predicted probability of a given peptide representing an epitope, and a measure of how unique to a given pathogen that potential epitope is. We may also wish to constrain our analysis to peptides that have a predicted probability greater than a minimal threshold, or that have some minimal amount of dissimilarity from the closest known epitope from another organism, to reduce the space of possibilities that a decision-maker will need to consider. These requirements give rise to the following problem formulation.

Let *𝒳* = {*x*_1_, *x*_2_, …, *x*_*N*_} represent the set of all candidate peptides returned by an epitope prediction model *M* for a given target pathogen, and let 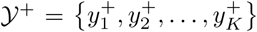 represent the set of known epitopes of all organisms except the target one. The multi-objective target selection problem can be defined as determining a subset *𝒳*\* ⊆ *𝒳* which is Pareto-optimal (see definition below) for the following formulation:

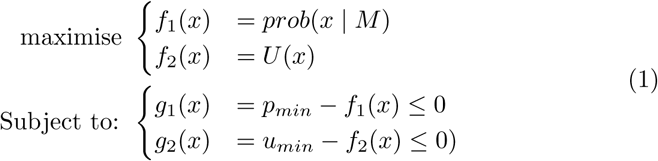

in which *prob*(*x* | *M*) denotes the probability score returned by model *M* for a candidate peptide *x*; *U* (*x*) is the uniqueness score of *x*, as defined in section “Uniqueness estimation” below; and *p*_*min*_ and *u*_*min*_ are user-defined thresholds for the smallest values of probability and dissimilarity to be considered in the prioritisation procedure. Notice that, since both *f*_1_(*x*) and *f*_2_(*x*) are strictly positive, the user can simply opt to set *p*_*min*_ = 0 and *u*_*min*_ = 0 to make the problem effectively unconstrained, if so desired.

Two definitions are required in order to understand what optimality means in the context of this problem.^1^ The first is the concept of *Pareto dominance* (Miettinen, 2012; Goulart and Campelo, 2016):

#### Definition 1 (Pareto dominance).

Given two alternative candidate solutions, *x*_1_ is said to *Pareto-dominate x*_2_ if and only if *f*_*i*_(*x*_1_) *≥ f*_*i*_(*x*_2_) for all *i*, with *f*_*i*_(*x*_1_) *> f*_*i*_(*x*_2_) for at least one *i*. This relationship is expressed as *x*_1_ ≻ *x*_2_ or, equivalently, as *f* (*x*_1_) ≻ *f* (*x*_2_). If neither alternative dominates the other, they are said to be incomparable or mutually non-dominated, a relation expressed by the operator ⊁.

The concept of Pareto dominance induces a partial ordering on the candidate solutions, meaning that some pairs have a clear ordering of preference, whereas others are incomparable - representing trade-offs of the objectives. Points that are not Pareto-dominated by any other in the set of feasible solutions are known as *Pareto-optimal* (Miettinen, 2012; Goulart and Campelo, 2016):

#### Definition 2 (Pareto optimality).

A candidate solution *x*\* is said to be *Pareto-optimal* (or *efficient*) if and only if ∄*x* ∈ *𝒳* | *x ≻ x*\*, i.e., if it is not dominated by any other feasible point. The set of all efficient points is known as the *efficient* or *Pareto-optimal set* of a problem, and its image in the space of objectives is called the *Pareto front*.

In general, multi-objective optimisation requires the use of iterative methods to explore the space of possible solutions and determine either the exact Pareto-optimal set (or a sample, in the case of continuous problems), or an approximation of this set. However, this is not needed for the the prioritisation problem considered in this work, as the finite and relatively small number of potential targets returned by any epitope prediction model makes it possible to estimate the uniqueness score of all candidates (as defined later) and rank all solutions by their dominance score, by simply executing a fast non-dominated sorting procedure on the full set of candidate peptides. The main concept of non-dominated sorting (Deb et al., 2002) is a simple procedure: (i) Given *𝒳*, compute all non-dominated elements and attribute the rank 1 to this set. (ii) Remove all rank-1 candidates, repeat the process and attribute the rank 2 to the new non-dominated set; and so on, until all points have received a dominance ranking. This fast procedure allows one to obtain a priority ranking of candidate peptides, based on their estimated probabilities of representing an epitope and their dissimilarity to known epitopes from other pathogens. The following sections describe how these two objectives are estimated in this work.

### Epitope prediction models (probability estimation)

To estimate both the candidate peptides and the values for the first objective, we need epitope predictors that are capable of returning a set of candidate peptides with associated probability scores. In this work we used organism-specific random forest predictors for three test pathogens.

These predictors were developed in an earlier work (Ashford et al., 2021), for the following pathogens: *Onchocerca volvulus*, Epstein-Barr Virus and Hepatitis C Virus. These organisms were selected due to the availability of reasonably balanced epitope data for them in the Immune Epitope Database (IEDB) (Vita et al., 2019). Data sets specific to each organism, based on taxonomy ID information, were generated from a full IEDB XML export. The data sets were filtered according to the criteria described in (Ashford et al., 2021) (Section 2.1, “Data Sets”). A fixed-width sliding window (length 15 amino acids) with a step size of one was applied over each sequence in the data sets to extract individual, fixed-length windows for feature calculation. The decisions behind the data representation and a list of all of the features calculated for each window can be found in our earlier work (Ashford et al., 2021) (Section 2.1.1, “Data Representation”). Prior to modelling, the organism-specific data sets were split into training and *Hold-out* data sets, with an approximate 75/25 split. To prevent potential data leakage (Kaufman et al., 2012), the data sets were split at the protein level based on both protein ID and local alignment scores. The training data sets were then used to develop random forest predictors specific to each organism. Information regarding the implementation of these predictors can also be found in our earlier work (Ashford et al., 2021) (Section 2.2, “Modelling”).

It is important to notice that, although we are employing the organism-specific models described above for the calculation of the estimated probabilities in the multi-objective formulation (1), any prediction model capable of outputting probabilities could be used instead. This includes most of the currently-used online predictors, such as BepiPred 2.0 (Jespersen et al., 2017), Lbtope (Singh et al., 2013), ABCpred (Saha and Raghava, 2006) or iBCE-EL (Manavalan et al., 2018).

### Sequence alignment (uniqueness estimation)

Amino-acid scoring matrices capture evolutionary relationships by quantifying differences that are likely to have occurred as lineages during evolutionary time. The least conservative changes receive the greatest negative scores, whereas conservative changes receive positive scores, resulting in values that are more informative than simple string dissimilarity measures. Pearson (Pearson, 2013) discusses the history of amino-acid substitution matrices, as well as their classification and penalty details, with a particular focus on BLOSUM and PAM scoring matrices. As the peptides that are compared in this work consist of relatively short amino-acid sequences, shallower matrices are considered preferable (Pearson, 2013), which resulted in the choice of PAM30 as the scoring matrix for all alignment calculations in this work.

Alignment scores were calculated using the *Biostrings* R library version 2.40.2 (Pagès et al., 2021), function *pairwiseAlignment*. The Smith-Waterman algorithm (Smith and Waterman, 1981) was used to obtain optimal alignment scores, with the gap opening penalty equal to 5 and the gap extension one set to 2.

Based on these definitions, the uniqueness score used for the multi-objective prioritisation was defined as follows. Let **S** *∈* ℝ^*N ×K*^ be a matrix containing the local alignment scores between the *N* candidate peptides in set *𝒳* and *K* known epitopes from organisms distinct from the target pathogen. Let also **a** *∈* ℝ^*N*^ be the vector of self-alignment scores of these *N* candidate peptides, and **b** *∈* ℝ^*K*^ be the vector of self-alignment scores of the *K* known epitopes. The uniqueness score of a candidate peptide *x*_*i*_ is calculated as:

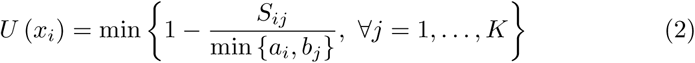

which returns a value between zero (perfect match between target *x*_*i*_ and a known epitope) and one (maximum dissimilarity given the length of *x*_*i*_). This uniqueness score is taken here as a proxy for a lower probability of crossreactivity during the development of the diagnostic test.

To obtain the set of known epitopes from non-target pathogens, the full set of positively labelled entries from the IEDB was extracted according to the same protocols described in our earlier work (Ashford et al., 2021). For each pathogen of interest, it was then filtered to remove all epitopes associated with that particular organism.

### Pipeline

The concepts introduced in the preceding sections are combined into the epitope prediction and target prioritisation pipeline illustrated in Figure 2. The probability calculations can be performed by any epitope prediction tool capable of outputting probabilities. Moreover, notice that the multi-objective approach presented in this work can be employed not only for prioritising linear B-cell epitope assessment, but also (by changing the uniqueness calculation function) for conformational epitope prediction/prioritisation or any other tasks in which multiple conflicting objectives may need to be considered simultaneously.

**Figure 2:**
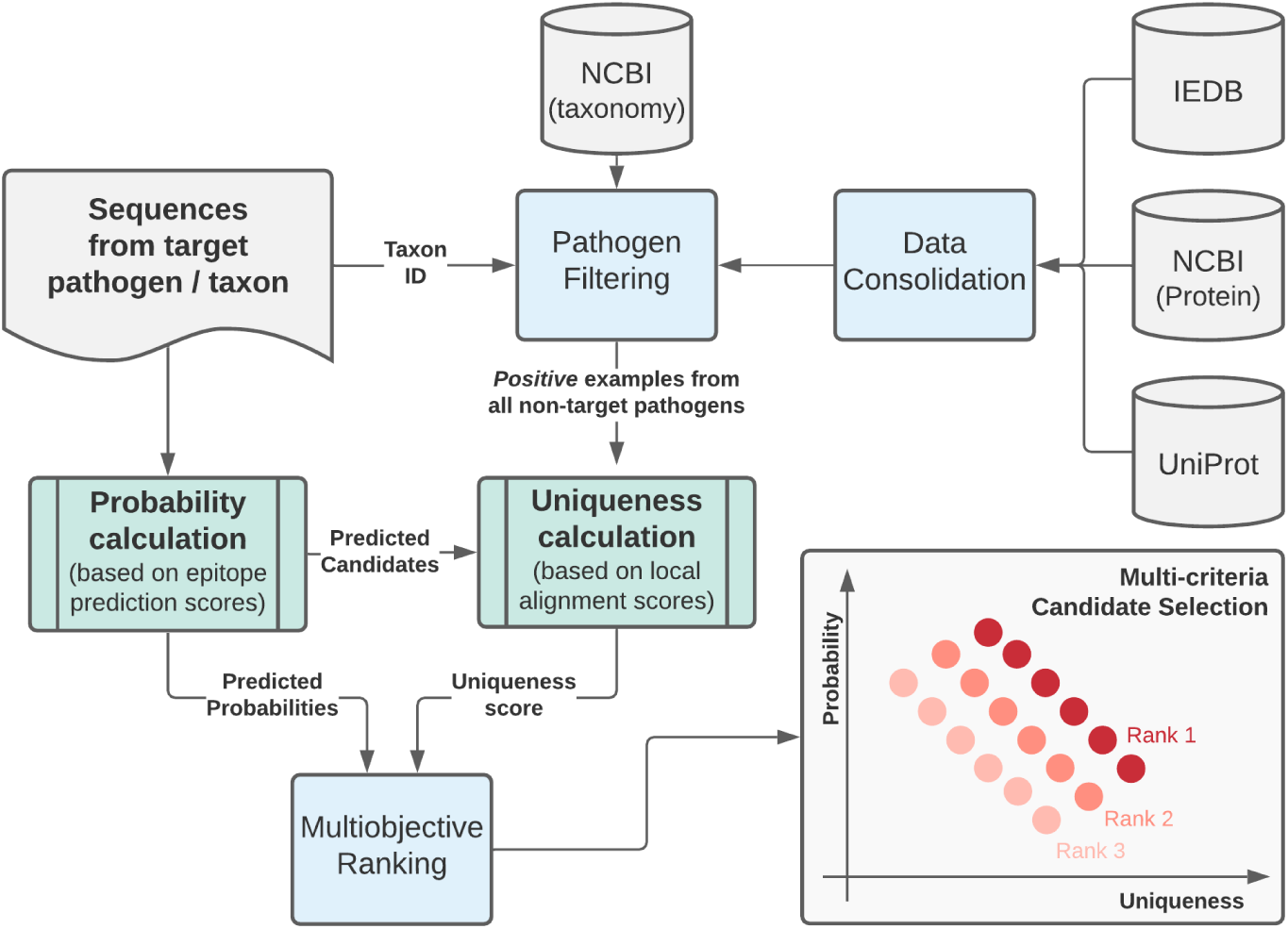
Multi-objective prediction and prioritisation pipeline. Publicly-available data on known epitopes (IEDB), protein sequences (NCBI-Protein and Uniprot) and organism taxonomy (NCBI-taxonomy) is used to extract all known epitopes or epitope-containing regions from organisms other than the target pathogen. An epitope predictor is used to extract candidate peptides from sequences of the target pathogen, associated with a probability score. The uniqueness score for each candidate peptide is calculated based on their smallest dissimilarity value to known epitopes from any organism other than the target pathogen. Non-dominated sorting is then used to derive priority rankings for each predicted peptide based on its probability and uniqueness values.

## Author’s contributions

FC defined the research question, developed the multi-objective model, designed the experiments and data visualisation. RC implemented the uniqueness calculations and non-dominated sorting routines, and ran the main experiments.

JA contributed the organism-specific predictions and literature review. JRC provided biological context, discussion and interpretation of the results. All authors contributed to and approved the final version of the manuscript.

## Acknowledgements

We would like to thank Dr. Francisco P. Lobo (UFMG, Brazil) for invaluable discussions that led to the selection of Smith-Waterman local alignment as the most adequate score for the uniqueness calculations.

## Footnotes

1 Notice that we are providing these definitions in the context of maximisation of all objective functions. Equivalent definitions can be easily derived for any combination of objective maximisation and minimisation.

## References

Ashford, J. ao Reis-Cunha, J., Lobo, I., Lobo, F., and Campelo, F. (2021). Organism-specific training improves performance of linear b-cell epitope prediction. Bioinformatics. Accepted for publication.

Blythe, M. J. and Flower, D. R. (2005). Benchmarking b cell epitope prediction: underperformance of existing methods. Protein Science, 14(1):246–248.

Can, H., Köseoğlu, A. E., Alak, S. E., Güvendi, M., Döşkaya, M., Karakavuk, M., Gürüz, A. Y., and ün, C. (2020). In silico discovery of antigenic proteins and epitopes of SARS-CoV-2 for the development of a vaccine or a diagnostic approach for COVID-19. Scientific Reports, 10(1).

Cancro, M. P. and Tomayko, M. M. (2021). Memory b cells and plasma cells: The differentiative continuum of humoral immunity. Immunological Reviews, 303(1):72–82.

Chen, J., Liu, H., Yang, J., and Chou, K.-C. (2007). Prediction of linear b-cell epitopes using amino acid pair antigenicity scale. Amino acids, 33(3):423–428.

Collatz, M., Mock, F., Barth, E., Hölzer, M., Sachse, K., and Marz, M. (2021). Epidope: A deep neural network for linear b-cell epitope prediction. Bioinformatics, 37(4):448–455.

Darricarrère, N., Pougatcheva, S., Duan, X., Rudicell, R. S., Chou, T.-H., DiNapoli, J., Ross, T. M., Alefantis, T., Vogel, T. U., Kleanthous, H., Wei, C.-J., and Nabel, G. J. (2018). Development of a pan-h1 influenza vaccine. Journal of Virology, 92(22).

de Oliveira Mendes, T. A., Cunha, J. L. R., de Almeida Lourdes, R., Luiz, G. F. R., Lemos, L. D., dos Santos, A. R. R., da Câmara, A. C. J., da Cunha Galvão, L. M., Bern, C., Gilman, R. H., Fujiwara, R. T., Gazzinelli, R. T., and Bartholomeu, D. C. (2013). Identification of strain-specific b-cell epitopes in trypanosoma cruzi using genome-scale epitope prediction and high-throughput immunoscreening with peptide arrays. PLoS Neglected Tropical Diseases, 7(10):e2524.

Deb, K., Pratap, A., Agarwal, S., and Meyarivan, T. (2002). A fast and elitist multiobjective genetic algorithm: Nsga-ii. IEEE transactions on evolutionary computation, 6(2):182–197.

EL-Manzalawy, Y., Dobbs, D., and Honavar, V. (2008). Predicting linear b-cell epitopes using string kernels. Journal of Molecular Recognition: An Interdisciplinary Journal, 21(4):243–255.

Gao, J., Faraggi, E., Zhou, Y., Ruan, J., and Kurgan, L. (2012). Best: Improved prediction of b-cell epitopes from antigen sequences. Plos One, 7(6):e40104.

Giacò, L., Amicosante, M., Fraziano, M., Gherardini, P. F., Ausiello, G., Helmer-Citterich, M., Colizzi, V., and Cabibbo, A. (2012). B-pred, a structure based b-cell epitopes prediction server. Advances and applications in bioinformatics and chemistry: AABC, 5:11.

Goulart, F. and Campelo, F. (2016). Preference-guided evolutionary algorithms for many-objective optimization. Information Sciences, 329:236–255. Special issue on Discovery Science.

Gourlay, L., Peri, C., Bolognesi, M., and Colombo, G. (2017). Structure and computation in immunoreagent design: From diagnostics to vaccines. Trends in Biotechnology, 35(12):1208–1220.

Handl, J., Kell, D. B., and Knowles, J. (2007). Multiobjective optimization in bioinformatics and computational biology. IEEE/ACM Transactions on Computational Biology and Bioinformatics, 4(2):279–292.

Haste Andersen, P., Nielsen, M., and Lund, O. (2006). Prediction of residues in discontinuous b-cell epitopes using protein 3d structures. Protein Science, 15(11):2558–2567.

Jespersen, M. C., Peters, B., Nielsen, M., and Marcatili, P. (2017). Bepipred-2.0: improving sequence-based b-cell epitope prediction using conformational epitopes. Nucleic acids research, 45(W1):W24–W29.

Kamath, K., Reifert, J., Johnston, T., Gable, C., Pantazes, R. J., Rivera, H. N., McAuliffe, I., Handali, S., and Daugherty, P. S. (2020). Antibody epitope repertoire analysis enables rapid antigen discovery and multiplex serology. Scientific reports, 10(1):1–9.

Kaufman, S., Rosset, S., Perlich, C., and Stitelman, O. (2012). Leakage in data mining: Formulation, detection, and avoidance. ACM Transactions on Knowledge Discovery from Data (TKDD), 6(4):1–21.

Lian, Y., Huang Zi, C., Ge, M., and Ming Pan, X. (2015). An improved method for predicting linear b-cell epitope using deep maxout networks. Biomedical and Environmental Sciences, 28(6).

Lin, S. Y.-H., Cheng, C.-W., and Su, E. C.-Y. (2013). Prediction of b-cell epitopes using evolutionary information and propensity scales. In BMC bioinformatics, volume 14, page S10. Springer.

Luo, Q., Qadri, F., Kansal, R., Rasko, D. A., Sheikh, A., and Fleckenstein, J. M. (2015). Conservation and immunogenicity of novel antigens in diverse isolates of enterotoxigenic escherichia coli. PLOS Neglected Tropical Diseases, 9(1):e0003446.

Manavalan, B., Govindaraj, R. G., Shin, T. H., Kim, M. O., and Lee, G. (2018). ibce-el: a new ensemble learning framework for improved linear b-cell epitope prediction. Frontiers in immunology, 9:1695.

Miettinen, K. (2012). Nonlinear multiobjective optimization, volume 12. Springer Science & Business Media.

Mosa, A. I. (2020). Antigenic variability. Frontiers in Immunology, 11.

Nascimento, I. and Leite, L. (2012). Recombinant vaccines and the development of new vaccine strategies. Brazilian journal of medical and biological research, 45:1102–1111.

Ong, E., Cooke, M. F., Huffman, A., Xiang, Z., Wong, M. U., Wang, H., Seetharaman, M., Valdez, N., and He, Y. (2021). Vaxign2: the second generation of the first web-based vaccine design program using reverse vaccinology and machine learning. Nucleic Acids Research, 49(W1):W671–W678.

Pagès, H., Aboyoun, P., Gentleman, R., and DebRoy, S. (2021). Biostrings: Efficient manipulation of biological strings. R package version 2.60.2.

Pearson, W. R. (2013). Selecting the right similarity-scoring matrix. Current protocols in bioinformatics, 43:3.5.1–3.5.9. 24509512[pmid].

Reis-Cunha, J. L., de Oliveira Mendes, T. A., de Almeida Lourdes, R., dos Santos Ribeiro, D. R., de Avila, R. A. M., de Oliveira Tavares, M., Lemos, D. S., Câmara, A. C. J., Olórtegui, C. C., de Lana, M., da Cunha Galvão, L. M., Fujiwara, R. T., and Bartholomeu, D. C. (2014). Genome-wide screening and identification of new trypanosoma cruzi antigens with potential application for chronic chagas disease diagnosis. PLoS ONE, 9(9):e106304.

Saha, S. and Raghava, G. P. S. (2006). Prediction of continuous b-cell epitopes in an antigen using recurrent neural network. Proteins: Structure, Function, and Bioinformatics, 65(1):40–48.

Sanchez-Trincado, J. L., Gomez-Perosanz, M., and Reche, P. A. (2017). Fundamentals and methods for t-and b-cell epitope prediction. Journal of immunology research, 2017.

Saravanan, V. and Gautham, N. (2015). Harnessing computational biology for exact linear b-cell epitope prediction: a novel amino acid composition-based feature descriptor. Omics: a journal of integrative biology, 19(10):648–658.

Shen, W., Cao, Y., Cha, L., Zhang, X., Ying, X., Zhang, W., Ge, K., Li, W., and Zhong, L. (2015). Predicting linear b-cell epitopes using amino acid anchoring pair composition. BioData mining, 8(1):1–12.

Sher, G., Zhi, D., and Zhang, S. (2017). Drrep: deep ridge regressed epitope predictor. BMC genomics, 18(6):55–65.

Shirai, H., Prades, C., Vita, R., Marcatili, P., Popovic, B., Xu, J., Overington, J. P., Hirayama, K., Soga, S., Tsunoyama, K., et al. (2014). Antibody informatics for drug discovery. Biochimica et Biophysica Acta (BBA)-Proteins and Proteomics, 1844(11):2002–2015.

Singh, H., Ansari, H. R., and Raghava, G. P. (2013). Improved method for linear b-cell epitope prediction using antigen’s primary sequence. PloS one, 8(5):e62216.

Smith, T. and Waterman, M. (1981). Identification of common molecular subsequences. Journal of Molecular Biology, 147(1):195–197.

Soria-Guerra, R. E., Nieto-Gomez, R., Govea-Alonso, D. O., and Rosales-Mendoza, S. (2015). An overview of bioinformatics tools for epitope prediction: implications on vaccine development. Journal of biomedical informatics, 53:405–414.

Suárez-Farinãs, M., Suprun, M., Kearney, P., Getts, R., Grishina, G., Hayward, C., Luta, D., Porter, A., Witmer, M., du Toit, G., et al. (2021). Accurate and reproducible diagnosis of peanut allergy using epitope mapping. Allergy.

Swenson, D. L., Warfield, K. L., Negley, D. L., Schmaljohn, A., Aman, M. J., and Bavari, S. (2005). Virus-like particles exhibit potential as a pan-filovirus vaccine for both ebola and marburg viral infections. Vaccine, 23(23):3033–3042.

Sweredoski, M. J. and Baldi, P. (2009). Cobepro: a novel system for predicting continuous b-cell epitopes. Protein Engineering, Design & Selection, 22(3):113–120.

Vita, R., Mahajan, S., Overton, J. A., Dhanda, S. K., Martini, S., Cantrell, J. R., Wheeler, D. K., Sette, A., and Peters, B. (2019). The immune epitope database (iedb): 2018 update. Nucleic acids research, 47(D1):D339–D343.

Wang, Y., Wu, W., Negre, N. N., White, K. P., Cheng, L., and Shah, P. K. (2011). Determinants of antigenicity and specificity in immune response for protein sequences. BMC Bioinformatics, 12:251.

Wee, L. J., Simarmata, D., Kam, Y.-W., Ng, L. F., and Tong, J. C. (2010). Svm-based prediction of linear b-cell epitopes using bayes feature extraction. In BMC genomics, volume 11, page S21. Springer.

Yang, X. and Yu, X. (2009). An introduction to epitope prediction methods and software. Reviews in medical virology, 19(2):77–96.

Yao, B., Zhang, L., Liang, S., and Zhang, C. (2012). Svmtrip: a method to predict antigenic epitopes using support vector machine to integrate tri-peptide similarity and propensity.

Yasser, E.-M. and Honavar, V. (2010). Recent advances in b-cell epitope prediction methods. Immunome research, 6(2):1–9.

Yasser, E.-M. and Honavar, V. (2014). Building classifier ensembles for b-cell epitope prediction. In Immunoinformatics, pages 285–294. Springer.

